# Improved synapsis dynamics accompany meiotic stability in *Arabidopsis arenosa* autotetraploids

**DOI:** 10.1101/2025.01.07.631194

**Authors:** Adrián Gonzalo, Aditya Nayak, Kirsten Bomblies

## Abstract

During meiosis, the correct pairing, synapsis, and recombination of homologous chromosome pairs is critical for fertility of sexual eukaryotes. These processes are challenged in polyploids, which possess additional copies of each chromosome. Polyploidy thus provides a unique context to study how evolution can modify meiotic programs in response to challenges. We previously observed that in *Arabidopsis arenosa* newly formed (neo-)polyploids, synapsis defects precede chromosomes associating in aberrant multivalent and univalent configurations. Here we study synapsis dynamics in genotypes with varying levels of meiotic stability to ask whether efficient synaptic progression is a key component of evolving stable tetraploid meiosis. We develop a method to quantify synapsis dynamics using the progression of foci of the pro-crossover factor HEI10 as a reference. HEI10 initially appears at many small loci before accumulating only at crossover sites. In diploids, this transition begins while there is still significant asynapsis, which quickly declines as HEI10 accumulation to fewer foci progresses. In neo-tetraploids, suboptimal elongation of synaptic initiation sites, and perhaps defective pairing, precedes synapsis stalling before the onset of HEI10 accumulation. In established tetraploids, HEI10 accumulation begins only when asynapsis is minimal, suggesting an enhanced HEI10/synapsis co-dynamic (even compared to diploids). Hybrids generated by crossing neo- and established tetraploids exhibit intermediate phenotypes. We find the extent of asynapsis correlates positively with crossover numbers, as well as a higher frequency of multivalents and univalent, which can disturb chromosome segregation. Our work supports the hypothesis that improving the efficiency of synapsis is important for evolving polyploid meiotic stability.

**Significance Statement:** During meiosis, homologous chromosomes pair and subsequently form the synaptonemal complex, which provides a context for maturation of DNA recombination events. How synapsis and crossover formation are coordinated in polyploids, where multiple chromosome sets complicate pair-wise interactions, remains unclear. Leveraging the dynamics of the pro-crossover factor HEI10 as a ‘developmental clock,’ we quantified synapsis progression in new and established tetraploids of *Arabidopsis arenosa*. Synapsis is severely compromised in neo-tetraploids, while in established ones it is more efficient even than diploids. Notably, the extent of synaptic defects correlated with excess crossovers, which compromises meiotic stability in tetraploids by connecting connect multiple homologs in multivalents. These findings highlight that improving synapsis dynamics are important for the evolution of meiotic stability in polyploids.

## Introduction

Meiosis is a specialized cell division that halves chromosome numbers through two rounds of segregation, producing haploid gametes. Crossovers generate new trait combinations and also hold homologous chromosomes in tight apposition, which ensures proper chromosome segregation – and thus fertility – in meiosis I (1). In polyploids, whole genome duplication leads to a genome with more than the usual two copies of each chromosomes, which can disrupt key meiotic processes, leading to abnormal chromosome configurations and impaired segregation (2). Understanding which processes are perturbed by genome duplication, and how they are adjusted in evolved polyploids, provides novel insights into meiosis and its evolution(3), as well as helping us understand how polyploids re-establish meiotic stability.

In diploids, proper pairing and segregation of homologs requires the correct number and pattern of crossovers, which is regulated by a meiotic program that ensures pair-wise interactions between homologs (forming bivalents) during prophase I of meiosis. Recombination begins with the formation of DNA double-strand breaks (4), which are processed in the context of linear proteinaceous axes that form along the length of each homolog (5). DSBs are typically required for subsequent homolog co-alignment, and polymerization of the synaptonemal complex during zygotene (6). The synaptonemal complex holds homologs in close alignment as recombination events mature, and its formation coincides with the partial removal of some axis proteins, such as ASY1 in plants (7, 8). The synaptonemal complex acts as a platform for recombination proteins, including the dosage-sensitive HEI10, which transitions from dispersed weak foci in early pachytene to bright foci at crossover sites in late pachytene (9–13). This process of HEI10 accumulation has been modeled based on data from *Arabidopsis thaliana* and *Caenorhabditis elegans* to follow the biophysics of a “coarsening” process that has been proposed to explain crossover patterning (14, 15). However, this model is debated, as it implies that crossover designation can occur later than proposed in well-supported models from budding yeast and *Sordaria* (16, 17). Regardless of whether HEI10 moves along it, defective synaptonemal complex formation is known to cause abnormal crossover numbers and patterns in some species, including *Arabidopsis* (18–21).

The processes of both synapsis and HEI10 accumulation are conserved across eukaryotic kingdoms (10, 11, 16, 22–24). However, it is not known how they might be affected by whole genome duplication or how they might evolve in response. In our previous work on natural auto-tetraploid *A. arenosa* (which arose from a single within-species whole genome duplication about 30,000 generations ago (25)) we found that defects in neo-polyploids in crossover patterning were accompanied by a defect in synapsis (26). Meiotically stable established tetraploids, in contrast, not only evolved improved crossover patterning, they also recovered full synapsis. It remained unknown at what stage synapsis becomes defective in neo-polyploids, whether it is stalled or merely slow, to what extent established tetraploid synapsis is enhanced, or whether the synaptic defects in neo-polyploids relate directly to defects in crossover patterning.

To clarify when and how synapsis becomes defective in neo-tetraploids and whether evolved tetraploids have more efficient synapsis, we studied the co-dynamics of synapsis and HEI10 progression. Quantifying these dynamics would ideally require live imaging (27), but this is limited to a few model species. Therefore, we developed a quantitative framework to assess synapsis dynamics in male meiocytes of four *A. arenosa* genotypes with varying meiotic stability. We found that synapsis is fully stalled, not just slow, in neo-tetraploids, which is associated with defects in chromosome co-alignment. Though synaptic initiation appears normal in the neo-tetraploids, they cannot efficiently elongate the synaptonemal complex from the initial sites. These defects correlate with increased crossover rates, suggesting crossover patterning issues, which in turn may cause the observed extensive multivalent and univalent formation. In contrast, established tetraploids exhibit more efficient synapsis, even surpassing diploids, suggesting that synapsis improvements contributed to meiotic stabilization and improved crossover regulation.

## Results

### Crossover number and meiotic stabilities of all genotypes at metaphase I

To explore the link between prophase I defects and meiotic instability in metaphase I, we quantified meiotic features in natural diploids (2X), neo-tetraploid lines, created by colchicine treatment in the lab (NEO-4X, see Material and Methods), evolved tetraploids (EST-4X), and hybrids (HYB-4X) from crosses between EST-4X and NEO-4X (Fig. S1). We analyzed metaphase I and diakinesis spreads (Fig. 1A, B). As previously reported, (26, 28), 2X showed full meiotic stability with all chromosome pairs forming bivalents. EST-4X formed on average less than 1 quadrivalent per cell (Fig. 1C) and very rarely univalents (0.1 per cell; Fig. 1D). HYB-4X had significantly more quadrivalents than EST-4X (Fig. 1C), and a similarly low number of univalents (Fig. 1D). In contrast, NEO-4X formed significantly more quadrivalents (3.0 ± 1.9 per cell, p < 0.0041, Fig. 1C) and also more univalents (0.9 ± 1.4 per cell, p > 0.0031, Fig. 1D).

**Figure 1.**
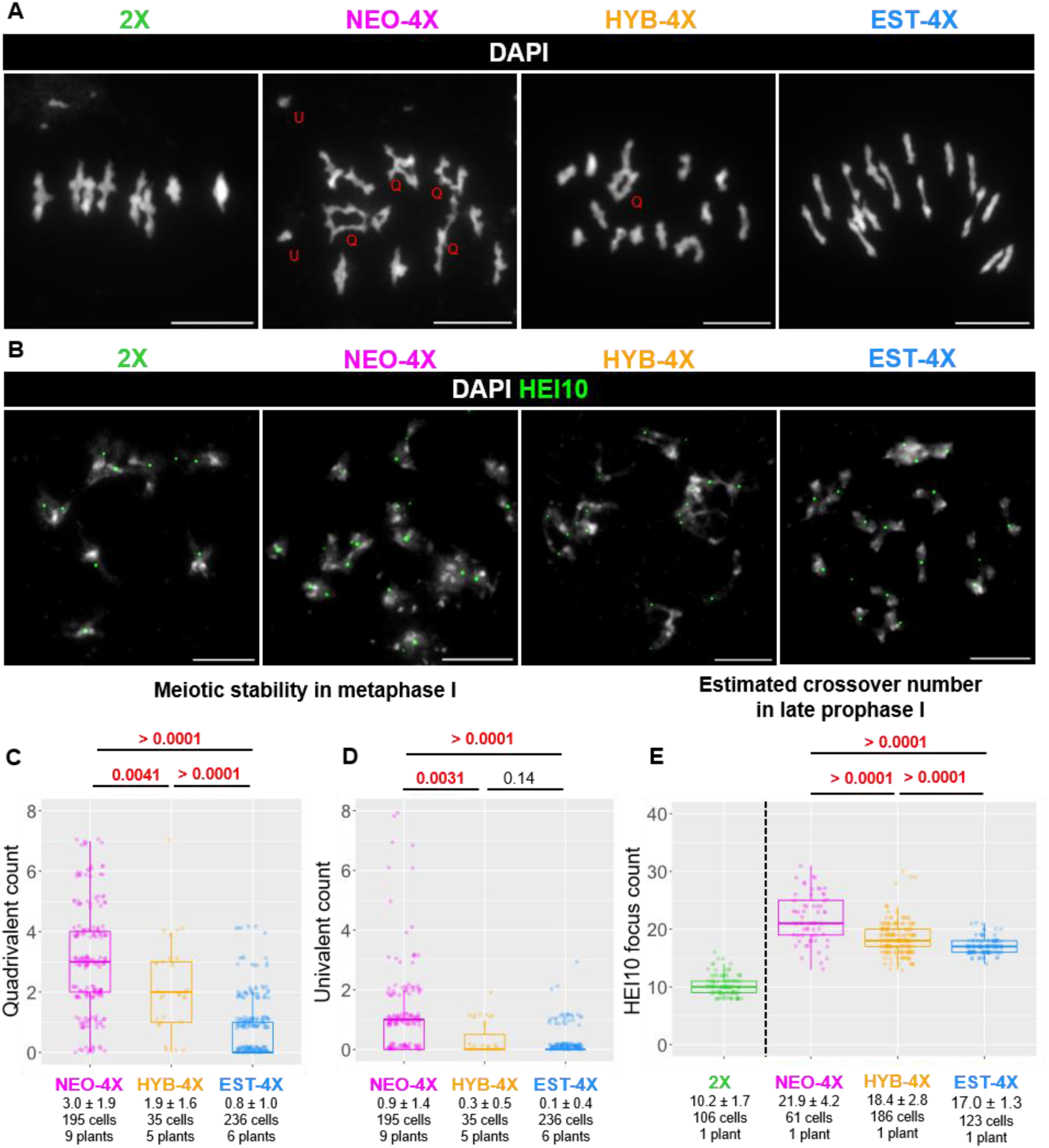
Crossover number and meiotic stability in different tetraploid populations. Scale Bar, 10 µm. (A) Examples of metaphase I spreads from all genotypes. Red Q, T, and U, mark examples of quadrivalents, trivalent, and univalent, respectively. (B) Examples of diakinesis cells with HEI10 immunostaining to detect crossovers. (C,D) Box plots of quadrivalent (C) and univalent (D) counts per cell. (E) HEI10 foci count in late prophase I (diplotene plus pachytene) cells. For (C), (D) and (E) and (E), population name, mean ± SD, and sample size. P-values are indicated on top of the plots for each comparison (according to Wald’s test on Negative binomial-GLMM, Poisson-GLMM coefficients for (C) and (D); and Dunnett’s test for (E).

Because the frequency of quadrivalents should be positively (and univalents negatively) correlated with crossover frequency (29–31), we also measured crossover number in 2X, NEO-4X, HYB-4X and EST-4X by immunostaining for HEI10, which in late prophase I stages of diplotene and diakinesis forms discrete foci at Class I crossover sites (32) (Fig. 1B). Consistent with previous reports (26) we scored ∼10 HEI10 foci per cell in 2X, about 22 in NEO-4X and 17 in EST-4X. HYB-4X had an intermediate number (about 18 foci) between its two parent genotypes (Fig. 1E), suggesting crossover number, like multivalent frequency, has a semidominant genetic basis.

### Quantification of HEI10 accumulation enables modeling of synapsis progression

To obtain a detailed picture of prophase I dynamics in the different tetraploid materials, we developed a quantitative framework by exploiting observable changes in HEI10 localization during pachytene as a “developmental clock.” HEI10 signal typically transitions from a very dispersed pattern of numerous weak foci in early pachytene, to a few bright and enlarged foci that mark the position of crossovers in late pachytene (10, 14, 22–24). We will refer to this dynamic process as “HEI10 accumulation.” We used HEI10 accumulation as a quantitative proxy for the developmental progression of meiosis to study the dynamics of other concomitant events, mainly synapsis.

We imaged male meiocytes in the prophase I stages of zygotene and pachytene in a set of 17 plants (Fig. S1), including 2X (4 plants), NEO-4X (6 plants), HYB-4X (3 plants), and EST-4X (4 plants) using Structured Illumination Microscopy (SIM). We immunolocalized three meiotic proteins (Fig. 2A): ZYP1 (the transverse filament of the synaptonemal complex, which marks synapsed axes), ASY1 (an axis component largely removed upon synapsis, thus serving as a marker for unsynapsed chromosomes), and HEI10 (to mark intermediates at different stages of the recombination process). Prior to image analysis, we verified that these three antibodies specifically stain meiotic cells; as expected, only a dim background signal was detected upon overexposure in somatic cells (Fig. S2A).

**Figure 2.**
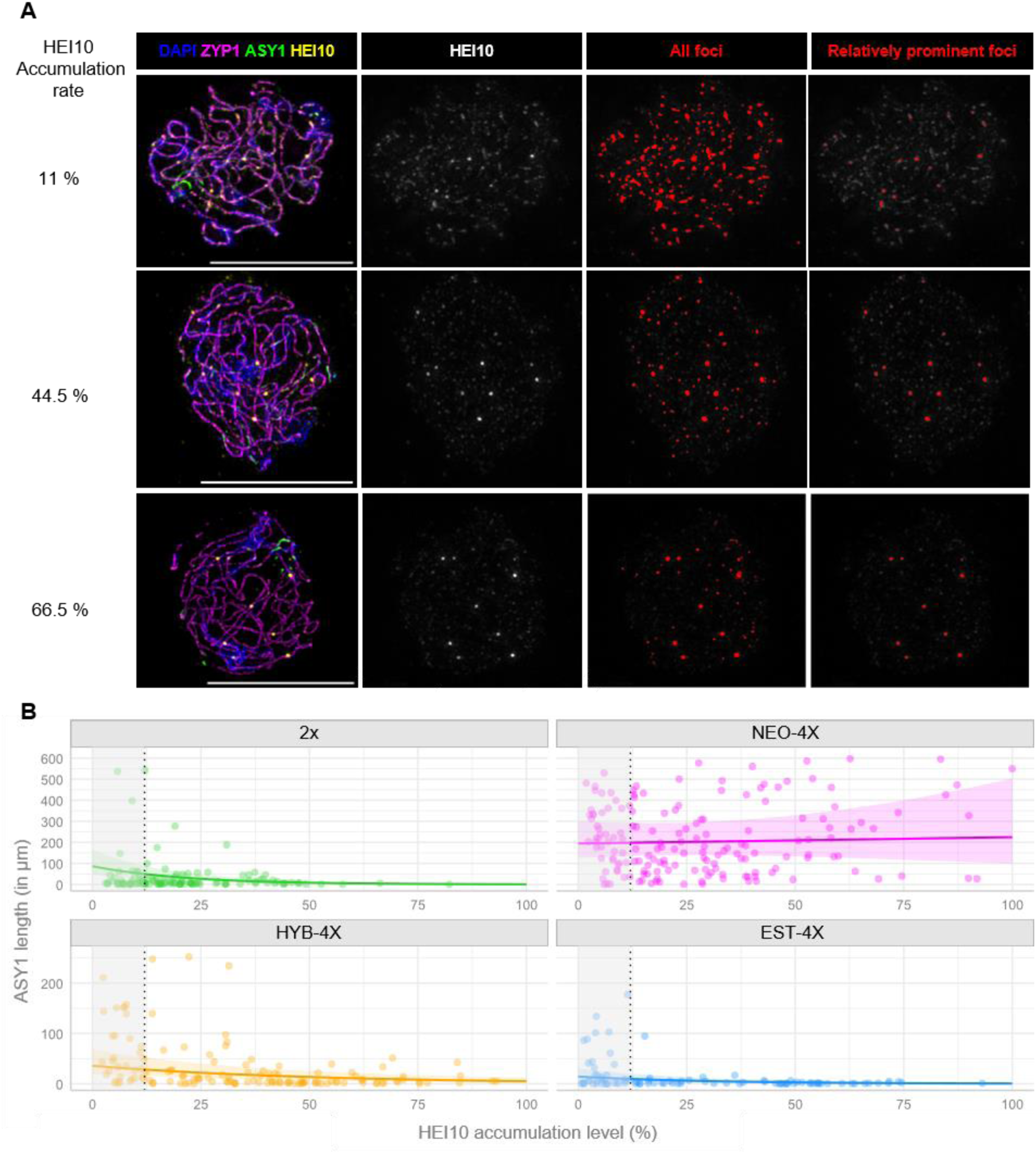
Modeling synapsis dynamics. Scale bar 10 microns. (A) Examples of imaged cells where two different thresholds (highlighted in red) were applied to the HEI10 channel to detect the total signal and prominent foci. Based on the percentage of signal in prominent foci compared to the total, the “HEI10 accumulation level” shown at left was calculated (see also Fig. S2). (B) Plots of all analyzed cells showing HEI10 accumulation level (x axis) by µm of ASY1 (asynapsis, in the Y-axis). Trendlines and 95% confidence intervals (shaded areas) are based in estimations from GLMM. Grey-shadowed region marks datapoints that were not included in GLMM analysis because they were under the HEI10 accumulation cutoff (12%) we estimated as our detection threshold.

Using a Fiji macro (see Material and Methods; Fig. 2A, S2B), we quantified HEI10 signal distribution while preventing bias from human decisions through particle analysis of HEI10 foci. The “HEI10 accumulation level” was calculated on each cell as the percentage of HEI10 signal intensity in the most relatively prominent foci compared to the total HEI10 signal intensity in that cell (Fig. 2A, Fig. S2B, C). We defined “prominent foci” vs. “total signal” using two different automated detection thresholds (Fig. 2A, S2B, C, see also Material and Methods). Each image serves as a “snapshot” of a dynamic process, showing diverse HEI10 accumulation levels across different cells (Fig. 2A, Fig. 3A), from low (where prominent foci account for a small share of total HEI10 signal intensity; Fig. 2A) to high (where most total HEI10 signal is concentrated in prominent foci, presumably marking crossover sites; Fig. 2A).

**Figure 3.**
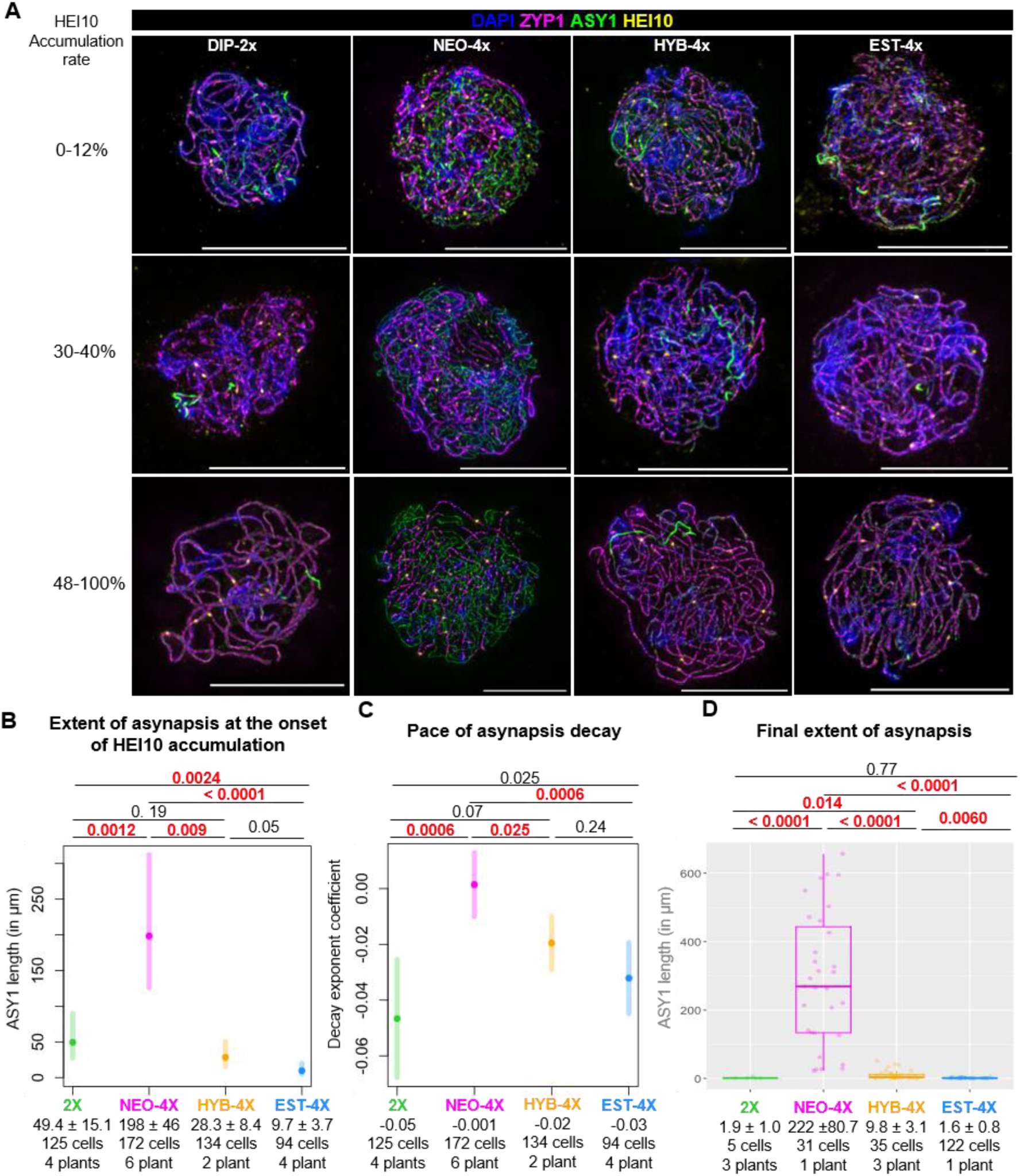
Differences in synapsis dynamics. Scale bar = 10 μm. (A) Examples of imaged cells of all genotypes with different levels of HEI10 accumulation showing how asynapsis (marked by ASY1, green) generally declines as HEI10 accumulation level increases, except in NEO-4X. (B) Plot of predicted extent of asynapsis by a Gamma-GLMM at the onset of HEI10 accumulation. Error bars represent 95% confidence intervals. (C) Plot of decay exponent coefficients estimated by a Gamma-GLMM for different genotypes. Error bars represent 95% confidence intervals. (D) is a box plot of the final level of asynapsis, at levels of HEI10 accumulation where synapsis does not progress further. P-values are indicated on top of the plots for each comparison (according to Wald’s test on Gamma-GLMM coefficients). Significant p-values were highlighted in red. Mean values (Mean ± SE) and sample sizes are indicated in the lower part of the plots.

To assess the potential effect of background signal on HEI10 accumulation, we measured somatic and meiotic leptotene cells, where HEI10 accumulation should be zero (Fig. S2D). These cells registered an average of 7% accumulation level, with a maximum of 12% (Fig. S2E). Therefore, we excluded cells with HEI10 accumulation levels below 12% from further analyses, as these values might be indistinguishable from background. This resulted in a final dataset of 523 imaged meiocytes from all genotypes (2X: 125 cells from 4 plants; NEO-4X: 172 cells from 6 plants; HYB-4X: 134 cells from 3 plants; EST-4X: 94 cells from 4 plants). To quantify synapsis levels, we used the ASY1 3D length (in µm) as a negative proxy for synapsis completion. Unfortunately, the punctate pattern of ZYP1 immunostaining precluded reliable automated measurement of its 3D length (Fig. S2F).

Next, we fitted a Gamma-Generalized Linear Mixed Model (Gamma-GLMM, see Material and Methods and SI Appendix) using HEI10 accumulation level and genotype as predictors (explanatory variables) for the extent of asynapsis as the response variable (Fig. 2B, Fig. S3A). GLMMs are especially recommended when data from different individuals (with different sample sizes each) are pooled to deal with pseudoreplication. This is particularly important in heterozygous outcrossing species like *A. arenosa*, which have extensive genetic and phenotypic variation among individuals (33). The individual plant is specified as a random effect in the model formula (see SI Appendix). This model suggests a negative correlation between HEI10 accumulation and extent of asynapsis, described by an exponential decay curve with substantial explanatory power (R² = 0.660; see Material and Methods and SI Appendix). This fits prior descriptions that large HEI10 foci marking crossovers are associated with full synapsis (10, 14, 22–24, 34). Overall, these analyses illustrate the utility of HEI10 accumulation level as a quantitative marker for developmental progression during mid-prophase I, and as a reference to quantify synapsis dynamics.

### Synapsis stalls in NEO-4X but is optimized in EST-4X

Despite the overall negative association between HEI10 accumulation level and the extent of asynapsis, plotting asynapsis (µm of ASY1 signal) against HEI10 accumulation revealed strong differences among genotypes, particularly for NEO-4X (Fig. 2B). Notably, NEO-4X exhibited higher levels of asynapsis at the onset of HEI10 accumulation (which we consider to be our above-calculated threshold of 12%) and little or no progress afterward, indicated by a flat trend line. To quantitatively analyze the differences in synapsis among genotypes, we examined the predictions and coefficients from our fitted model (Fig. 3B, C, D). We doubled the values of ASY1 length in 2X so they can be directly compared with those of tetraploids. At the onset of HEI10 accumulation (defined as our 12% cutoff), the extent of asynapsis was notably higher in NEO-4X (nearly 200 µm of ASY1; Fig. 3A, B) compared to 2X (about 50 µm of ASY1). In contrast, EST-4X showed minor asynapsis at this point (less than 10 µm of ASY1), significantly different from both NEO-4X and 2X (Fig. 3A, B). HYB-4X presented intermediate values (almost 30 µm ASY1; Fig. 3A, B), not significantly different from 2X or EST-4X but differing from NEO-4X. Overall, this analysis demonstrates that asynapsis at the onset of HEI10 accumulation varies significantly across genotypes, with NEO-4X exhibiting the highest levels. Interestingly, EST-4X displayed lower asynapsis than 2X, indicating it has more efficient early synapsis relative to HEI10 accumulation.

The analyses above raised the question of whether synapsis in NEO-4X is terminally stalled or merely very slow. Our Gamma-GLMM allowed us to compare the pace of synapsis progression across all genotypes by testing for differences in decay exponent coefficients of ASY1 length relative to HEI10 accumulation levels (Fig. 3C). We found statistically comparable decay exponent coefficients in 2X, HYB-4X, and EST-4X (Fig. 3D), which are all significantly different from 0 (p < 0.0001 for all). These means that ASY1 length changes exponentially as HEI10 accumulation increases. In contrast, the decay exponent of NEO-4X was not significantly different from 0 (p = 0.8, Fig. 3A and 3D) and differed significantly from all other genotypes (Fig. 3D), —hence the flat trendline observed in Fig. 2B. This result suggests that NEO-4X does not significantly reduce asynapsis as HEI10 accumulation advances, indicating that the extensive asynapsis present at the beginning persists through to the end of pachytene. These findings support the hypothesis that NEO-4X synapsis stalls before HEI10 accumulation begins. Although GLMM analysis excels in dealing with differences between individuals, plotting each plant individually hinds some differences in behavior (Fig S3), so it should not be ruled out that some individual plants may differ from the overall trend.

We also examined the final levels of asynapsis by determining when asynapsis ceases to decline. To do so, we identified the HEI10 accumulation level at which decay exponent coefficients for all genotypes were not significantly different from zero. We found that models including only cells with HEI10 accumulation levels greater than 49% had decay exponent coefficients for all genotypes not significantly different from zero (at least, p = 0.12, Gamma-GLMM, R² = 0.849). This model suggests, that after this cutoff, cells have already reached their asynapsis minimum. At this HEI10 accumulation level, both 2X and EST-4X had reached full synapsis (less than 2 µm of ASY1, Fig. 3D, S4). NEO-4X in contrast, retained very high levels of asynapsis (Fig. 3D, S4) and interestinlgy HYB-4X showed intermediate levels (almost 10 of µm of ASY1) significantly different from both all the other genotypes.

### Neo-tetraploids have less co-alignment of late asynaptic regions

Having observed that synapsis stalls in NEO-4X already before HEI10 starts accumulating, we hypothesized asynapsis in NEO-4X may be due to even earlier problems, e.g., during pairing, which leads to co-alignment. Thus, we examined co-alignment in cells with 30-150 µm of linear ASY1 signals, which allows assessment relative axis position (this becomes difficult in cells with greater ASY1 lengths). In this sub-sampled set we observed two types of geometry: “irregular”, where pairs of asynaptic axes are not near one another, and “parallel”, where asynaptic axes could be easily observed as coaligned ASY1 signals (Fig. 4A,B). These parallel asynaptic structures were typically 0.5 to 10 µm long, and could be seen as either terminal asynaptic regions at chromosome ends, or interstitial bubbles where asynaptic regions lie between two synapsed regions (Fig. 4A,B). We believe these parallel asynapsis regions are regions that did not have time to synapse yet (or in NEO-4X had stalled) as opposed to unresolved interlocks, as we did not observe interlocks when these structures were examined in 3D. We scored the number of parallel and irregular structures per cell and used it as a response variable in a Negative-binomial-GLMM (Fig. S5A) where ASY1 length was the explanatory variable. We indeed observed (with a substantial explanatory power; R^2^= 0.7) that there was a positive correlation between the length of ASY1 and the number of parallel axes. We interrogated this model to ask whether there are differences in the number of parallel axes between genotypes at comparable levels of asynapsis (ASY1 length = 100 microns), and found that NEO-4X has fewer parallel asynaptic axes than the other three genotypes (at most p < 0.0052, Negative-binomial-GLMM, Fig. 4C).

**Figure 4.**
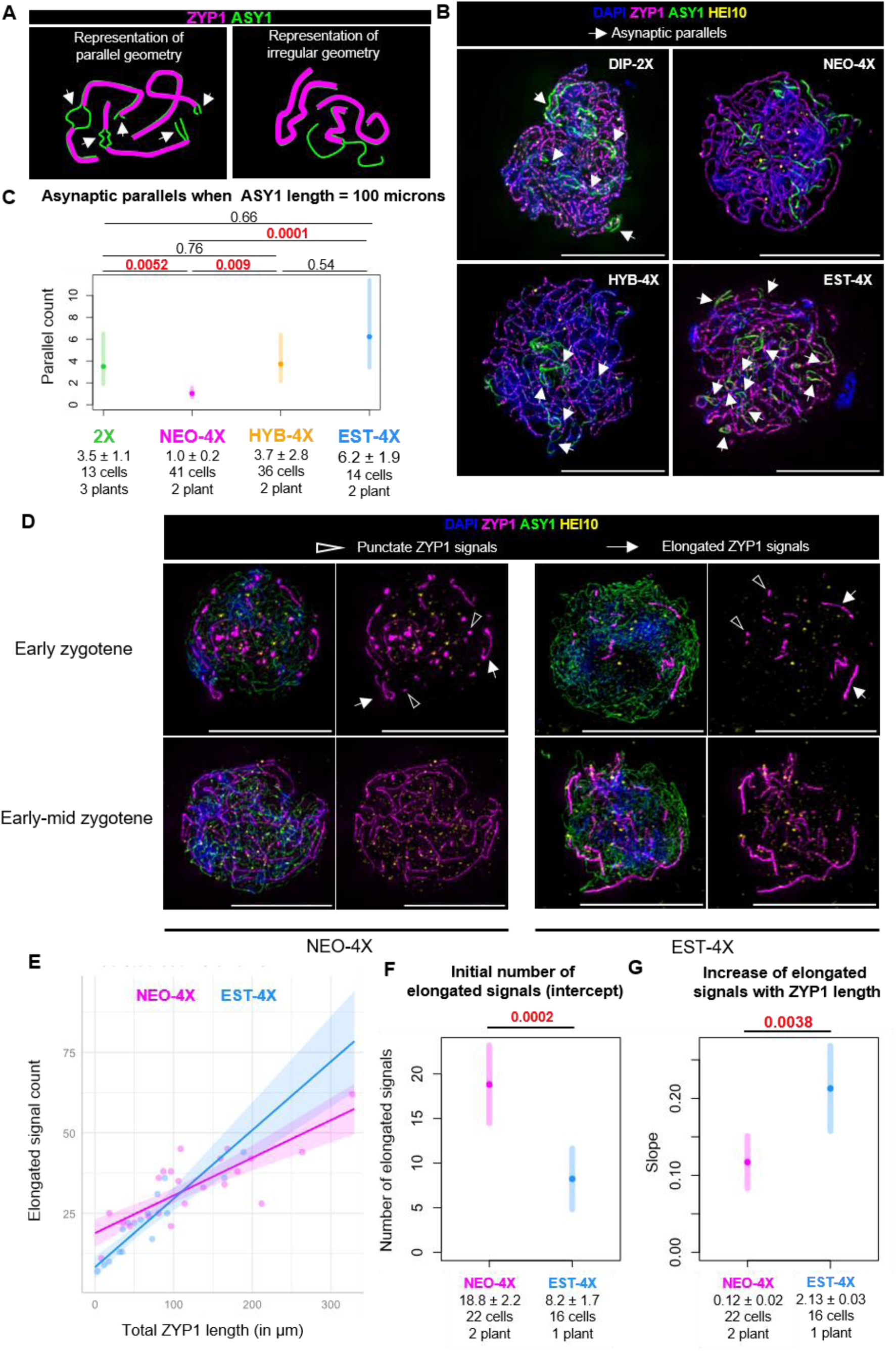
Early synaptic defects in NEO-4X. Scale bar = 10 µm. (A) Example illustration of two types of geometry of asynaptic regions observed: parallel and irregular. (B) Examples of these structures observed in cells from the four genotypes. Arrowheads indicate regions classed as “parallel”. (C) GLMM results showing differences in number of parallels when ASY1 length is 100 µm in all the genotypes, indicating differences in the frequencies of these structures. (D) Examples of early and early-to-mid zygotene meiocytes showing punctate (shaded arrowheads) and elongated (white arrowheads) ZYP1 signals. (E) Plot of data and predicted trendlines for NEO-4X and EST-4X according to our fitted model (Poisson-GLM) for elongated ZYP1 segments by total ZYP1 length. (F,G) Plots showing differences predicted by the fitted model in (F) the intercept for both genotypes (the initial number of SES, when ZYP1 length is zero) and (G) the slope (the increase of the number of SES as ZYP1 length grows). P-values are indicated on top of the plots for each comparison (according to Wald’s test on Poisson-GLMM coefficients). Significant p-values were highlighted in red. Mean values (Mean ± SE) and sample sizes are indicated in the lower part of the plots.

### EST-4X and NEO-4X show different early synapsis dynamics

Issues with pairing might also manifest as problems with synaptic initiation and/or elongation (19, 35), so we sought to get more insights into the initiation of synapsis by examining ZYP1 signals in early to mid-zygotene cells with little synapsis. We observed two kinds of ZYP1 signals in these cells: punctate signals, likely corresponding to synapsis initiation sites (SIS), and more elongated signals that we interpret as events where synapsis has begun to elongate from a previous synapsis initiation site (synapsis elongation sites; SES). To better understand the dynamics of ZYP1 signals, we first modeled their number (response variable) as a function of total ZYP1 length (explanatory variable). Although we cannot rule out that some synaptic stretches are reverted, we used the total length of ZYP1 as a proxy of developmental time (the longer the total ZYP1 length, the later the progress). Here, we used Generalized Linear Models (GLM, which differs from GLMM in that it does not control for individual variation or any other random effect) as they showed a better fit to the data than GLMMs (see SI Appendix).

Although we observed no differences in the behavior of punctate signals (Fig S5B, C and D), when we modeled the dynamics of the appearance of elongated ones, we fitted a negative-binomial-GLM with very strong explanatory power (R^2^=0.98, Fig. 4E), which suggests a linear correlation between the number of SES and the total length of ZYP1. Our GLM analysis showed a greater initial number of SES (the model’s intercept) in NEO-4X than EST-4X (Fig. 4F). Interestingly, we also observed that the rate at which the number of elongated signals increases as the total ZYP1 length grows (the model’s slope) is significantly greater in EST-4X than in NEO-4X. Since we already saw NEO-4X has fewer co-aligned regions and does not complete synapsis, we hypothesize that these differences in slope mean that synapsis initiation sites elongate more readily in EST-4X, whereas in NEO-4X synapsis initiates normally, but a lower proportion of the initiation sites are proficient to elongate normally.

### Preferential localization of prominent HEI10 foci to synapsed regions is stage-dependent

We next asked whether or how HEI10 focus development is affected by the synaptic defects in NEO-4X by examining HEI10 localization in more detail. It has been shown in other species that HEI10 forms foci at synaptic initiation sites (11, 17, 36–38), thus, we began by analyzing cells in early zygotene, when synapsis is just starting. Irrespective of genotype, in nuclei with incipient synapsis, the few prominent HEI10 foci localize with no apparent preference to either asynaptic axis (which accounts for the vast majority of axis at this stage) or occasionally on the few small ZYP1 signals where synapsis has initiated (early zygotene cells in Fig 4D). This observation is consistent with reports in *Arabidopsis thaliana* at the earliest stages of synapsis (10, 39), but not other species, where HEI10 orthologs are specifically associated with synapsis initiation sites (11, 17, 36, 37). Although, non-prominent foci are more difficult to distinguish from background signal, the lack of preference of prominent foci for synapsed stretches, suggests that in *A. arenosa*, HEI10 only marks (at most) a subset of synapsis initiation sites, and is not a reliable marker for them. In all genotypes, HEI10 is mostly found in discrete punctate foci, though on rare occasions, we observed linear HEI10 signal on unsynapsed chromosomes overlapping with ASY1, resembling a pattern previously described in wheat (24) (Fig S6). Importantly, we did not consider these cells in the previously described analyses as these linear signals could skew the results.

In more advanced stages, where HEI10 condensation to fewer foci has begun (HEI10 accumulation levels of 12-24%), we observed that prominent HEI10 foci are more commonly located on synapsed regions in all genotypes (early-mid zygotene cells in Fig. 4D, 5A, S6A,B), though they can occasionally also occur on unsynapsed regions at this stage. Later, at 45-100% HEI10 accumulation levels, enlarged discrete foci were found almost exclusively in synapsed regions. This was true for 2X, HYB-4X, EST-4X, that are fully or nearly fully synapsed, but also in NEO-4X, that still retains extensive asynapsis where only 3 of 919 prominent foci were localized on unsynapsed axis (Fig 5A). Remarkably, we observed that in these late asynaptic NEO-4X cells, even small tracts of synapsis (< 5 µm) can contain a prominent HEI10 focus (Fig 5B,C), resembling observations in mutants with patchy synapsis in *C. elegans* (18, 20) and *A. thaliana* (21, 39).

**Figure 5.**
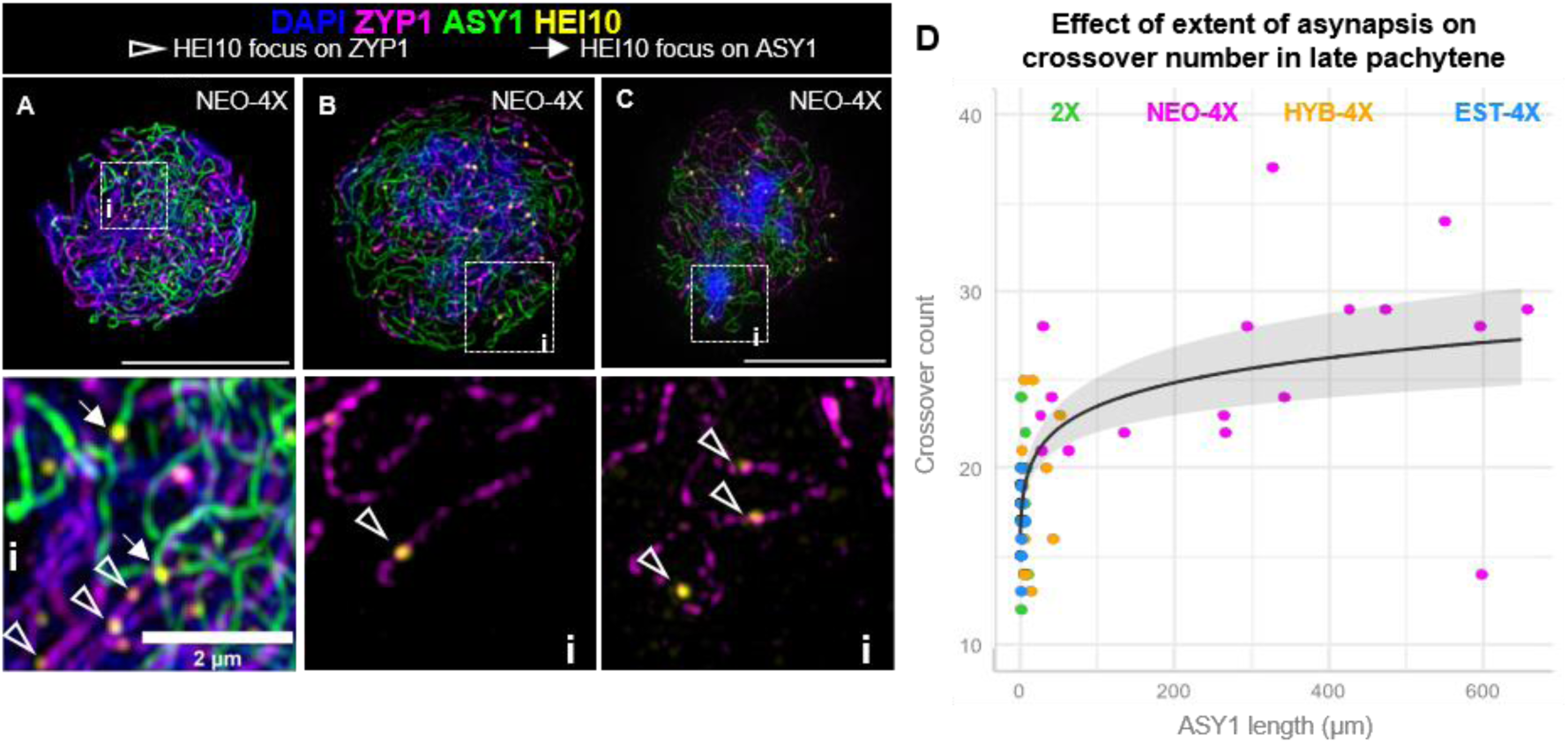
Relationship of synapsis with HEI10 localization and crossover number. (A) An example of a mid-zygotene cell in NEO-4X with prominent HEI10 foci co-localizing with both asynaptic (ASY1) and synapsed (ZYP1) axes. (Ai) Zoomed-in image of the dashed square in (A). (B) and (C), and their respective zoom in details (Bi) and (Ci) show examples of late pachytene-like cells with persisting asynapsis where prominent late HEI10 foci (presumably marking crossovers) specifically localize to synapsed chromosomes (marked by ZYP1). Notably, even short ZYP1 stretches can have late HEI10 foci. (D) Plot showing the exponential relationship between extent of asynapsis and crossover number for late pachytene(-like) cells.

### Extensive asynapsis accompanies elevated crossover number

Since synaptic alterations have been shown to alter crossover number and/or patterning in other species (18–21), we asked whether defective synapsis in NEO-4X is related to their elevated crossover number. Since the three tetraploid genotypes (NEO-4X, HYB-4X, EST-4X) have different levels of asynapsis (Fig. 3) and different crossover numbers (Fig. 1), we tested if there is a correlation among cells within a cytotype between degree of asynapsis and crossover number. We filtered for cells where HEI10 foci likely specifically mark crossovers (late pachytene), by determining after what HEI10 accumulation level the number of detected prominent foci resembled the most the crossover count at diakinesis for that genotype. This corresponded to HEI10 accumulation levels greater than 44, 59, 48, and 65% for 2X, NEO-4X, HYB-4X, and EST-4X, respectively (Fig. S7B). Once we calculated the number of HEI10 foci in each late pachytene cell, we used it as a response variable in a Poisson-GLM with the extent of asynapsis (µm of ASY1) as an explanatory variable. Our Poisson-GLM suggested, with substantial explanatory power (R^2^=0.608) that there is a significant (p < 0.0001) logarithmic relationship between the extent of asynapsis and crossover number (Fig. 5D). This non-linear relationship, along with the fact that we found no correlation within genotypes (R^2^=0.141, at most), suggests that the potential relationship between asynapsis and mis-regulation of crossovers is likely complex.

## Discussion

We previously showed that appropriate crossover number and patterning can prevent meiotic instability in autopolyploids (26, 29). We previously also reported that the synaptonemal complex is compromised in neo-tetraploids of *Arabidopsis arenosa* (26). Synaptic defects have also been reported in neo-autotetraploid *C. elegans* (40), as well as neo-allopolyploid *A. suecica* (41). Interestingly, established tetraploids of both *A. suecica* and *A. arenosa* exhibit complete synapsis (26, 42), suggesting its re-establishment is important for polyploids. In this study, we thus explored the dynamics of synapsis in both neo- and established tetraploid *A. arenosa* to better understand the origin of the problem itself, and more importantly, to ask if improving the dynamics of this process might be an important adaptation for meiotic stability in established tetraploids.

Prophase I, the stage where crossover patterning and synapsis take place (6), is a highly dynamic stage of meiosis. Quantitative analysis of a dynamic process like synapsis ideally would involve live imaging experiments (27), but this limits image resolution, and is not tenable in non-model systems where appropriate fluorescently tagged lines are not available. Moreover, tagging HEI10 at least in the *C. elegans* ortholog ZHP-3, has been reported to compromise its wild type function (43). An appealing alternative is to measure quantifiable dynamics of one process relative to quantifiable dynamics of another process in images of fixed meiocytes, which also allows analysis using super-resolution microscopy. Such an approach has been used to study e.g., the dynamics of DSB repair relative to synapsis progression in *A. thaliana* (44). Here we developed this approach to analyze synapsis dynamics using HEI10 progression as a reference (Fig. 6). While informative, this approach does carry the caveat that it provides only relative information and does not measure how long events take in real time.

Recent work in several species has focused on the role that the synaptonemal complex (which links the axes of homologous chromosomes and provides a platform for the maturation and regulation of recombination (45–48)), plays in the final number and pattern of crossovers on chromosomes (47, 49–51). Comparing synaptic progression to the progression of HEI10 accumulation, we found that there are differences among genotypes in synapsis dynamics already from the very beginning of the process (Fig. 6). Our data suggest that synapsis initiation occurs seemingly normally in neo-polyploids but elongates less efficiently in NEO-4X than EST-4X. This may result from defects in homologous axis pairing or synaptonemal complex polymerization. The former is supported based on asynaptic axes being more commonly parallel in 2X, HYB-4X, and EST-4X than in NEO-4X. In late zygotene in NEO-4X, many regions remain unaligned, likely unpaired, which may directly hinder synapsis completion. This diverges somewhat from the conclusions in autotetraploid *C. elegans*, where synaptic defects were observed even when axes were co-aligned in groups of four, leading to the idea that homologous chromosomes may “compete for establishing synapsis connections” (40). The same may happen locally in *A. arenosa* as well, where regional co-alignment of all four axes has also been observed, but these four-way alignments occur in neo-as well as established polyploids, and thus they seem in themselves to not block synapsis (26). It is nevertheless certainly possible that there are pairing problems that arise from competition among the four homologs in neo-polyploids that could hinder synaptonemal complex elongation, while the established polyploids evolved a solution to this challenge. Another explanation for inefficient synapsis in neo-polyploids, could be that extra chromosome sets complicate chromosome movements during prophase I, which are important to promote pairing and synapsis (19, 35, 52).

How might evolved tetraploids have solved the synaptic issues their neo-tetraploid counterparts face? Previously, we identified genes that show evidence of having been targets of natural selection in the established tetraploid *A. arenosa* lineage, including numerous potentially relevant meiosis genes (28, 53–55). If pairing itself is the ultimate problem, a possible candidate for the evolved solution in established tetraploids is *PRD3* (56), which shows strong evidence of having been targeted by selection in tetraploid *A. arenosa* (28). *PRD3* encodes a homolog of yeast *MER2* (57), and is essential for the formation of the DSBs required for pairing and synapsis (56, 58). Evolved variants of *PRD3* could thus enhance pairing, but whether or how they might do so remains to be tested. DNA strand invasion is a requisite in plants for pairing and synapsis (59, 60) which is regulated antagonistically by SDS (56) and FLIP(61) (and other factors), and the genes encoding them also show strong evidence of selection in the tetraploid lineage (55). *RMF1*, another gene under with evidence of having been targeted by selection in *A. arenosa*, was also recently shown to negatively regulate strand invasion (62). We also previously suggested that changes to the chromosome axis (e.g., stiffening), could lead to more efficient chromosome pairing and synapsis in the established tetraploids (26, 33). This which would fit with the strong evidence for selection having targeted the axis proteins ASY1 and ASY3 in the tetraploid lineage (28, 55), and where tetraploid alleles have already been shown to reduce multivalent frequency. Moreover, recently ASY3 dosage has been shown to affect synapsis and crossover number in allotetraploid *Brassica napus* (63). However, genetically generated established tetraploid plants homozygous for diploid alleles of ASY1 and ASY3 do not have defects in synapsis, suggesting that while they may contribute, they are not solely responsible for the difference in synapsis between diploids and tetraploid genotypes (33). Prominently, selection also targeted the synaptonemal complex protein ZYP1 (28, 55), suggesting the derived allele found in tetraploids may have a very direct role of modifying the synaptonemal complex itself, perhaps by enhancing the efficiency of polymerization, but this, too, remains to be tested.

An important question regarding the synaptic defects observed in neo-polyploids is whether they relate directly to the increased recombination rate observed in neo-tetraploids of *A. arenosa* (26) and other species (64, 65). We first found a broad correlation across genotypes with neo-tetraploids having both defective synapsis as well as higher crossover numbers relative to both established tetraploids and neotetraploid/established tetraploid hybrids. We found additional support for this correlation from the finding that asynapsis and crossover number correlate positively across cells, which showed a non-linear but logarithmic correlation. However, this non-linear relationship could indicate that crossover number is not proportional to the extent of asynapsis, but maybe to other factors like the number of synaptic patches (as observed in A. thaliana *pch2* mutants (21)). Alternatively, a threshold-like effect could disrupt crossover regulation at certain level of asynapsis, but not less. For example, DSBs are known to be increased in situations where homologs fail to engage through synapsis or crossovers in budding yeast and *C. elegans* (46, 66, 67).

We hypothesize this link may be caused at least in part by the fact that neo-tetraploids have HEI10 dynamics taking place in the context of abnormal patchy synapsis. This might allow recombination intermediates that would otherwise not develop into crossovers to accumulate sufficient HEI10 to become crossover-fated. Similar alterations of crossover patterning are reported for the *axr1* (39), *pss1* (19)and *pch2* (21) mutants in *A. thaliana* and *syp-1* knock down mutants in C*. elegans* (18, 20) which have patchy synapsis. Because of its preference for associating with synapsed regions, HEI10 accumulates locally to higher concentrations per length of synaptonemal complex when synapsis is patchy, resulting in nearly every synaptic “patch” developing at least one prominent HEI10 focus (19). Importantly, this would imply that local concentration of HEI10 in synapsed regions can alter the fate of at least some recombination intermediates, implying they retain some plasticity and remain sensitive to HEI10 levels.

We suggest that patchy synapsis in neo-tetraploids might similarly locally concentrate HEI10, assuming about the same amount of protein per unit DNA now accumulates on a smaller total amount of synaptonemal complex length. Conversely, in the context of full synapsis (as in the established tetraploids), HEI10 could be siphoned off recombination intermediates that are not irreversibly crossover-fated; in the context of patchy synapsis this may not be possible, allowing HEI10 to be retained on these plastic sites. The HEI10 present on a synaptic patch would accumulate where it can – a crossover-designated site when one is available, a plastic site when not. When this happens in *A. thaliana* patchy synapsis mutants, it does not result in a crossover increase, but this is because the number of synaptic patches is lower than the wildtype crossover number. However, in *C. elegans* where in wild type meiosis there is only 1 crossover per bivalent, mutants with patchy synapsis do show a crossover increase (18, 20). In *A. arenosa* crossover rates per chromosome pair are also low (1.06 and 1.25 crossovers per chromosome pair in EST-4X and 2X, respectively), and the number of synaptic patches in neo-tetraploids is greater than this (26). Overall, the increased efficiency of synapsis we see in the evolved polyploids of *A. arenosa* likely helps prevent excess recombination intermediates taking on crossover fates, and may also help solve other patterning issues (26).

## Materials and Methods

### Plant material

We used lab propagated descendants of natural populations of diploid (2X) and autotetraploid (EST-4X) *A. arenosa* from Strečno, Slovakia (SNO, 2X) and TBG Triberg, Germany (TBG, EST-4X) (25, 68). To produce neo-tetraploids (NEO-4X), we treated 2X 14-days-old seedlings in the apical meristem with 0.05% colchicine (Sigma) diluted in sterile water with 0.05% Silwet-77 (Anawa). Neo-tetraploid branches of treated plants were identified in colchicine-derived chimeric plants using flow cytometry. Confirmed tetraploid branches were used both for crosses (to obtain a second-generation non-chimeric neo-tetraploids). For the experiments we used both colchicine-treated and second-generation neo-tetraploids (as indicated in Figure S1), since our analysis showed no significant differences between them (see SI Appendix). To generate tetraploid hybrids (HYB-4X), we crossed EST-4X as male with NEO-4X as a female (as described in Figure S1). We verified the karyotype of second-generation NEO-4X, HYB-4X and EST-4X plants in meiotic and mitotic spreads and only proceed to further analyses with confirmed euploids.

### Metaphase I and late prophase I spreads

We used chromosomes spreads of flower buds fixed in ethanol:acetic acid 3:1 at diplotene, diakinesis (immunostaining of HEI10 for crossover quantification), and metaphase I (univalent and quadrivalent counts). We performed spreads following the protocol in ref. (32), with the minor modifications explained in ref. (29). Spreads were visualized and imaged using Leica Thunder Imager 3D Tissue epifluorescence microscope.

### Immunostaining

For HEI10 foci count for crossover quantification in late prophase I cells (diplotene and diakinesis) we followed the protocol in ref. (32), with the same minor modifications explained in ref. (29). We used a primary monoclonal antibody against *A. thaliana* HEI10 (29) and a secondary goat anti-guinea pig Alexa-488 antibody (both at 1:200 dilution). We visualized and imaged immunostained late prophase I cells with a Leica Thunder Imager 3D Tissue epifluorescence microscope. To visualize HEI10 accumulation and the extent of asynapsis in zygotene/pachytene cells, we used 3D Structured Illumination Microscopy (SIM) following the protocol described in ref. (69). For this immunostaining protocol, we used the following primary antibodies: rabbit polyclonal anti-HEI10, guinea pig monoclonal anti-ZYP1, guinea pig polyclonal anti-ZYP1, guinea pig polyclonal anti-ASY1 (all with 1:500 dilution) and rat polyclonal anti-ASY1 (with 1:1000 dilution). We used the following secondary antibodies with a 1:200 dilution: goat anti-rabbit Alexa-647, goat anti-rat Alexa-555, goat anti-guinea pig Alexa-488. Cells were imaged using a Deltavision OMX SIM microscope in 3D stacks of 0.125 microns optical section spacing.

## Data Generation and Analysis

### Image analysis and scoring

Multivalent and univalent scoring was performed blindly in randomized metaphase I images. Each metaphase I image was assigned to a “scorability” class (A, B, C, D, or E), depending on the quality of the spread and the confidence of the count, with A the best and E the least reliable (examples shown in Fig. S8). Scorability class was considered for statistical analyses. 2-chanel multi-image files for HEI10 focus count in late prophase I spreads did not allow image randomization. To analyzed HEI10 accumulation and the extent of asynapsis in zygotene/pachytene cells, we processed SIM images in 3D stacks using three different ImageJ Fiji macros applied in batch to the full image set. These macros apply different thresholding methods to detect fluorescence signals (70, 71) of HEI10 or ASY1. Macro 1 analyzes HEI10 signal to estimate the number of prominent foci, and the intensity of both prominent foci and total signal (which we used to calculate the HEI10 accumulation level). Macro 2, analyzes ASY1 signal to measure its 3D length (in µm) to estimate the extent of asynapsis. Macro 3 analyzes the signal from both ASY1 and ZYP1 for quality control purposes. Details about the macros are further explained in the SI Appendix.

### Statistical analysis

All statistical analyses were performed using Rstudio with the version 4.4.0 of R, and plots were made using ggplot2 (72). To analyze late prophase I HEI10 focus count data, since only one plant per individual was included per genotype, we used the classical statistical tests. Since there was unequal variance, we used Kruskal-Wallis and Dunnett tests (for multiple group comparisons) or t-test/Mann-Whitney U for simple comparisons (depending on whether equal variance criteria is met or not, respectively; details in Supplementary Dataset S8). For other datasets, since there were always genotypes with more than one individual, we used Generalized Linear Mixed Models, to account for random variation between individuals and prevent pseudoreplication. Occasionally we used Generalized Linear Models (which do not control for variability between individuals), when they showed a better fit than GLMMs (see SI Appendix). We used gaussian-GLM(M) or Gamma-GLM(M) (more appropriate for skewed data) for continuous data (such as ASY1 length) and Poisson-GLMM or negative binomial-GLMM for discrete data (such as multivalent/univalent or foci counts). We used the glmmTMB R package (73) to fit models. For each analysis we fitted several models and chose the best fit using the Performance (74) and DHARMa (75) R packages. Details on criteria and procedure to select the best fit are explained in the SI Appendix and Supporting Dataset S8 and S9. For the best fit GLMM, we estimated means, confidence intervals, medians, standard errors and p-values using the emmeans R package (76) with Holm-Bonferroni correction, based on Wald’s tests performed by glmmTMB.

### Data, Materials, and Software Availability

All image data used in this study are available freely from the ETH Research Collection. For SIM immunostaining images from 2x, DOI: 10.3929/ethz-b-000696798. For SIM immunostaining images from NEO-4X, DOI: 10.3929/ethz-b-000696907. For SIM immunostaining images from HYB-4X, DOI: 10.3929/ethz-b-000696850. For SIM immunostaining images from EST-4X, DOI: 10.3929/ethz-b-000696797. For epifluorescence microscopy images in metaphase, diakinesis and diplotene spreads, DOI; 10.3929/ethz-b-000696798. The ImageJ Fiji macros, R scripts and raw data files to reproduce the statistical analyses are available in https://github.com/adgon/MeioScope.

## Acknowledgments

We thank the entire Bomblies group for scientific discussions, particularly Marinela Dukic for constructive feedback and Tadeas Priklopil for critical reading and guidance on statistics. We are thankful to Sophie Thüring for technical support in preparing metaphase I spreads. We also thank Joiselle B. Fernandes for scientific discussions, critical reading of the manuscript and support in editing. We also thank Tobias Schwarz and ETH ScopeM for technical support with SIM. We thank Chris Franklin for providing some of the antibodies used in this work. This project has received funding from the European Union’s Horizon 2020 research and innovation programme under the Marie Sklodowska-Curie (MSC) grant agreement No 101029732 to AG, as well as ETH core funds (KB).

## Author Contributions

A.G. and K.B. conceptualized the research, A.G. and A.N. obtained the materials and reagents, A.G. performed the research. A.G. analyzed the data, A.G. and K.B. wrote and edited the manuscript.

## Competing Interest Statement

The authors declare no competing interests.

## Classification

Biological Sciences. Genetics.

## Notes

https://doi.org/10.3929/ethz-b-000696796

https://doi.org/10.3929/ethz-b-000696907

https://doi.org/10.3929/ethz-b-000696850

https://doi.org/10.3929/ethz-b-000696797

https://doi.org/10.3929/ethz-b-000696798

https://github.com/adgon/MeioScope

### Competing Interest Statement

The authors have declared no competing interest.

### Summary of Updates

Figure 5 was not included in the originally submitted file. This version does include Figure 5 and its corresponding legend.

