## Supplementary information for "Improved synapsis dynamics accompany meiotic stability in *Arabidopsis arenosa* autotetraploids"

**This PDF file includes:**

Supporting text
Figures S1 to S9
Table S1
Legends for Movies S1 to S9
SI References

**Other supporting materials for this manuscript include the following:**

Datasets S1 to S9

### Supporting Information Text

**General procedure for automated analysis of SIM images**. Each SIM image stack was firstly inspected. Stacks with images having too much background signal in one of the channels were discarded. Moreover, image stacks that lacked one of the channels were also discarded since the Fiji macros used only work on images with four channels. The macros are designed to be applied in files with channels ordered as follows: DAPI, ASY1, ZYP1, HEI10. files with a different order of channels need to be reordered. This can be done adding one initial step to the existing macro using the "Arrange channels" command. Moreover, these macros require that the images contain scale information in their metadata that can be accessible by Fiji. All macros are available in https://github.com/adgon/MeioScope.

Each macro, processes the signal from one or more channel in two steps. The first step is signal detection and the criterion of what is detected and signal and what is not is determined by different preestablished thresholding methods, namely; Yen(1)), Moments (2) and default (ordered from less to more sensitivity). The second step is the measurements on the detected signals which can be, signal count, intensity measurement, or length measurements.

Macro 1 analyzes HEI10 channel and provides two different output tables. In the first of them, with the count data of prominent foci (detected with Yen) that that totally or partially overlap with synapsed regions (ZYP1 signal, detected with default). The second output provides the HEI10 intensities from both prominent foci (detected with Yen) and the total signal (detected with Moments). The percentage of intensity from prominent foci relative to the total signal was later used to calculate the HEI10 accumulation level. For this analysis each and every cell was processed with automatic thresholding, allowing no exception for manual thresholding to prevent any bias.

Macro 2 measures the total 3D of ASY1 whose signal was thresholded using Yen. This measurement provides a measure of the extent of asynapsis. In the vast majority of cells, the threshold detects ASY1 highly specifically at asynaptic regions where the axis is not remodeled(3) and intense ASY1 signals persist, while the dim signal at remodeled axis after synapsis are not detected (Fig. S2F). However, in some rare cells, manual thresholding was required (Fig. S9).

Macro 3 (optional), quantifies 2D length of ASY1 and ZYP1 linear signals (thresholded with the Yen and default methods, respectively). These data were not analyzed in this work, but they can be useful for quality control purposes, to identify cells where manual thresholding is required. For instance, rare cells with an excessive length of both ASY1 and ZYP1, correspond to cases where ASY1 was overestimated (Fig. S9). Conversely, cases where ASY1 length in 2D is greater than in 3D, indicates an underestimation. Finally, visual inspection ASY1 thresholds can also be used for total certainty.

**General procedure for statistical modeling**. Generalized Linear Mixed Models (GLMMs) are used to predict the behavior of a response variable (e.g., ASY1 length or univalent count) as a function of one or more explanatory variables. Explanatory variables of primary interest, which are assumed to be measured without error, are referred to as fixed effects (e.g., HEI10 accumulation level or genotype). Explanatory variables that are not of primary interest but need to be controlled for, and are assumed to be drawn from a larger population, are referred to as random effects (e.g., individual or scorability class). The formulas for both fixed and random effects are specified in Supporting Dataset S9. In our models, we considered either a single random effect (e.g. plant individual) or a nested random effect (e.g., individual nested in genotype). Random effects models with more than two variables, or complex nested structures, were too complex given the amount of data available. When random effects were not included, we used Generalized Linear Models (GLMs) instead. To predict the behavior of a response variable using GLMs or GLMMS, it is necessary to specify the distribution family that reflects the nature of the response variable. For continuous variables (such as ASY1 length) we used Gamma or Gaussian distributions, whereas for discrete count data (such as multivalent counts), we used Poisson or negative binomial distributions. The negative binomial distribution was selected for count data when overdispersion (i.e., variance greater than the mean) was observed, as it accounts for the extra variability in the data. Additionally, generalized models require a link function, which transforms the linear combination of explanatory variables to the scale of the response variable and determines the shape of the predicted curve. For example, the log link function confers an exponential relationship between the predictors and the response. In cases where the data contained an excess of zero values, we included a zero-inflation term to appropriately handle the overabundance of zeros in the data set.

In each analysis, several models were fitted using different formulas (varying random effects, distributions, link functions, etc.; see Supporting Dataset S9), with the glmmTMB (4) R package used for all cases except for negative binomial-GLMs, which were fitted using the MASS R package. Models that failed to fit the data when using glmmTMB were discarded. Next, we used the Performance R package (5) to compare model metrics. Specifically, we compared the coefficients of determination ($R^2$) across models. For GLMMs, two types of $R^2$ were calculated: the conditional $R^2$, which accounts for both fixed and random effects, and the marginal $R^2$, which considers only fixed effects. In linear models, $R^2$ indicates the proportion of observed variability that can be explained by the model (larger $R^2$ suggesting a better fit)  It should be noticed, however, that the interpretation of the $R^2$ is more complex GLM(M)s although this metric it still used as it has similar properties in terms of explanatory power. (see (5) for detailed interpretation in the context of GLM/GLMMs). Additionally, we evaluated models using Akaike's Information Criterion (AIC) and Bayesian Information Criterion (BIC), which measure the balance between model accuracy and complexity.  Lower AIC and BIC values indicate better-performing models, with these criteria helping to penalize overfitting while still prioritizing explanatory power. Based on these metrics (detailed for each model fitted in Supporting Dataset S9), we selected the most promising models for each analysis for further residuals-based model diagnostic checks.

For residuals-based model diagnostics, we used the DHARMa R package (6). First, we verified that the variables included in the model did not exhibit collinearity. Additionally, we ensured that the model was not "singular", which is a strong indication of either collinearity or an overly complex and overfitted model (6). Models detected as singular were discarded due to their potential to have near-zero power. Next, we used the residuals to test four key assumptions: normality of residuals, absence of over- or under-dispersion, no excess of outliers, and homogeneity of variance (detailed descriptions of these tests are provided in Supporting Dataset S9). For testing variance homogeneity, DHARMa conducts multiple tests and generates diagnostic plots of the residuals. In cases where these plots showed no clear patterns and variance was roughly homogeneous across groups (e.g. genotypes), we tolerated minor heteroskedasticity (i.e., variance heterogeneity of the residuals). To select the best fitting model for each analysis, we prioritized models that had the best Performance metrics ($R^2$, AIC, and BIC) and succesfully passed the DHARMa diagnostics. Further details for each analysis are provided below.

**Modeling multivalent count in Metaphase I**. We tested different random effect structures, considering "genotype", "plant", and "scorability" classes as random effects. As explained in the Data Generation and Analysis section, the scorability class reflects the level of confidence in scoring each cell. However, because models including all three random effects failed the singularity test, we proceeded with models containing only one or two random effects terms. Models with a single random effect term performed better, with "scorability" class outperforming individual and nested structures involving both terms. Additionally, models that included only cells from the best scorability classes (A and B) performed better than those that incorporated all cellsclasses. Therefore, for further analysis, we compared models that used only these high-quality cells.

**Modeling univalent count in Metaphase I**. The modelling of univalent count data followed a similar approach as described for multivalent count (see above), with a difference that this time, a nested structure of random effects (involving both "scorability" class and "plant" individual) provided a better performance.

**Modeling asynapsis decay with HEI10 accumulation**. Minor heteroskedasticity (heterogeneous variance) was an issue in all the models tested, but the best fitting model did not show any discernible pattern in the residuals plot. Notably, variance was homogeneous both across and within genotypes.

**Modeling late asynapsis decay with HEI10 accumulation**. For this analysis, there were no significant issues to report.

**Modeling the effect of colchicine on NEO-4X behavior**. For consistency, we used the same model formula applied to the analysis of asynapsis decay with HEI10 accumulation. No additional models were evaluated. The selected model had minor heteroskedasticity, though no obvious patterns were present in the residual plots.

**Modeling dynamics of synapsis initiation**. All models that included random effects were singular, indicating that the model complexity exceeded what the data could support. Therefore, only GLMs were tested. Although count data typically requires discrete distributions, Gaussian models showed reasonable performance, which warranted testing. However, the best-fitting model was ultimately based on count-data distributions (Poisson).

**Modeling dynamics of early synapsis elongation.** This analysis the exhibited the same issues as those encountered during the modelling of synapsis initiation.

**Modeling crossover increase with asynapsis**. Applying a log transformation to the explanatory variable substantially improved the models, suggesting that the relationship was logarithmical rather than linear. In addition, GLMs performed better than GLMMs which, in all cases, were singular and with lower explanatory power.

**Modeling the effect of extensive asynapsis on crossover number**. Most models tested showed signs of overfitting (singularity, but not collinearity). The best fitting model without singularity did not include random effects and was fit using GLM. While minor heteroskedasticity was present, the variance was homogeneous across both plants and genotypes.

**Modeling decay of asynaptic parallels with ASY1 length**. For this analysis, there were no significant issues to report.

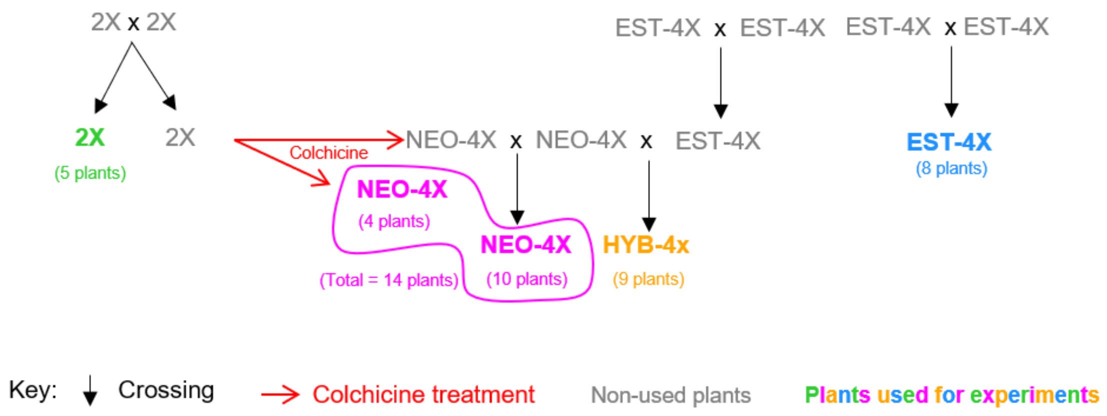


**Fig. S1.** Scheme of plant materials generated and used in this study.

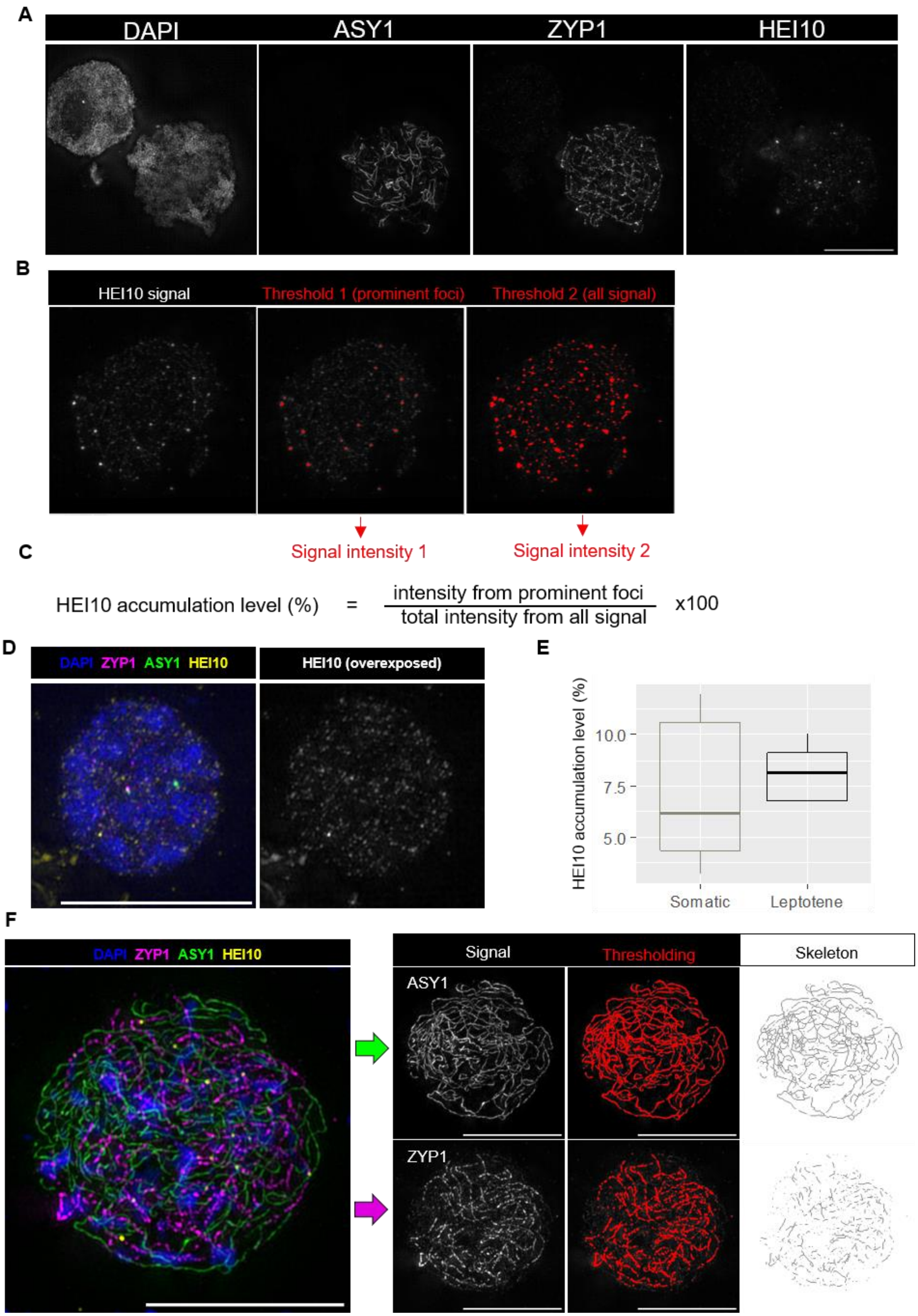
A
DAPI
ASY1
ZYP1
HEI10
B
HEI10 signal
Threshold 1 (prominent foci)
Threshold 2 (all signal)
Signal intensity 1
Signal intensity 2
C
HEI10 accumulation level (%) = intensity from prominent foci / total intensity from all signal x100
D
DAPI ZYP1 ASY1 HEI10
HEI10 (overexposed)
E
HEI10 accumulation level (%)
10.0
7.5
5.0
Somatic
Leptotene
F
DAPI ZYP1 ASY1 HEI10
Signal
Thresholding
Skeleton
ASY1
ZYP1

**Fig. S2.** Quantification of HEI10 accumulation level and extent of asynapsis. (A) Fluorescence staining for two cells: a somatic (only DAPI stains it substantially) with a meiotic cell (with specific staining for DAPI, ASY1, ZYP1 and HEI10) at the lower right. (B) Example images showing the HEI10 channel of one imaged cell showing the two thresholds assigned in Fiji: Threshold 1 (Yen method (1)) detects the relatively prominent foci, whereas Threshold 2 (Moments method (2)) detects the total HEI10 signal. (C) How HEI10 accumulation level is calculated from the values of signal intensity from Threshold 1 and Threshold 2. (D) The background signal from HEI10 staining in a somatic cell (signal has been overexposed for display). (E) The HEI10 accumulation values of leptotene and somatic cells (n=11, calculated in NEO-4X cells) where no HEI10 loading is expected. (F) Illustrates how we quantify ASY1 and ZYP1 length a NEO-4X cells with extensive asynapsis in Fiji. Each channel is processed separately. After thresholding, detected signal is skeletonized and measured. This example illustrates how ASY1 signal generates good quality skeletons whereas ZYP1 signal, which often displays a dotty pattern, yields discontinuous skeletons which severely underestimates its real length in automated measurements.

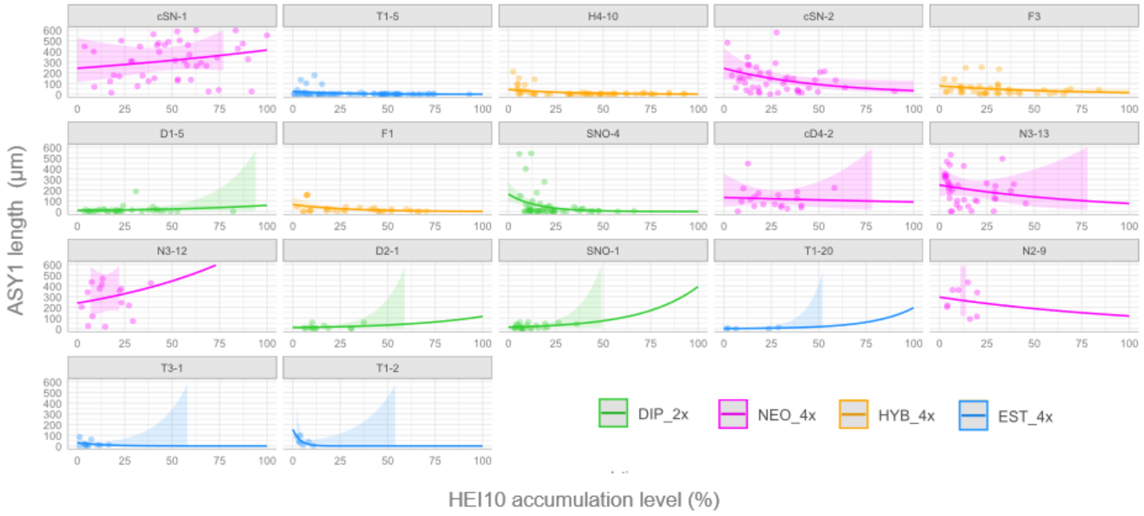


**Fig. S3** Synaptic behavior of multiple individuals.

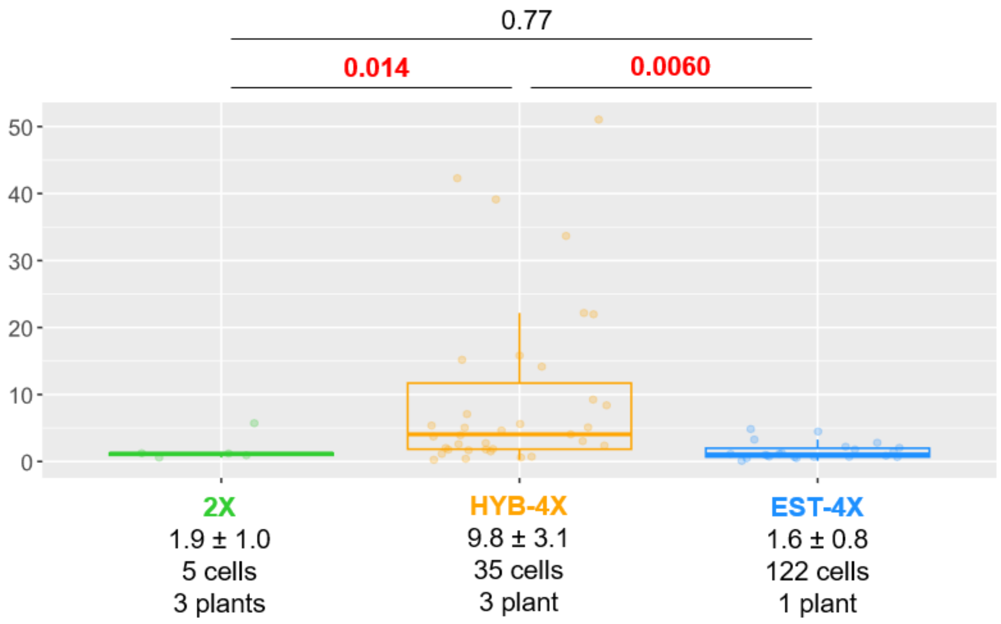


Fig. S4. Final levels of asynapsis. This is an alternative to the one in figure 3D, without NEO-4X in order to have a scale that allows better visualization of the data of the genotypes with very low levels of asynapsis. P-values are indicated on top of the plots for each comparison (according to Wald's test on Gamma-GLMM coefficients). Significant p- values were highlighted in red. Mean values (Mean ± SE) and sample sizes are indicated in the lower part of the plots.

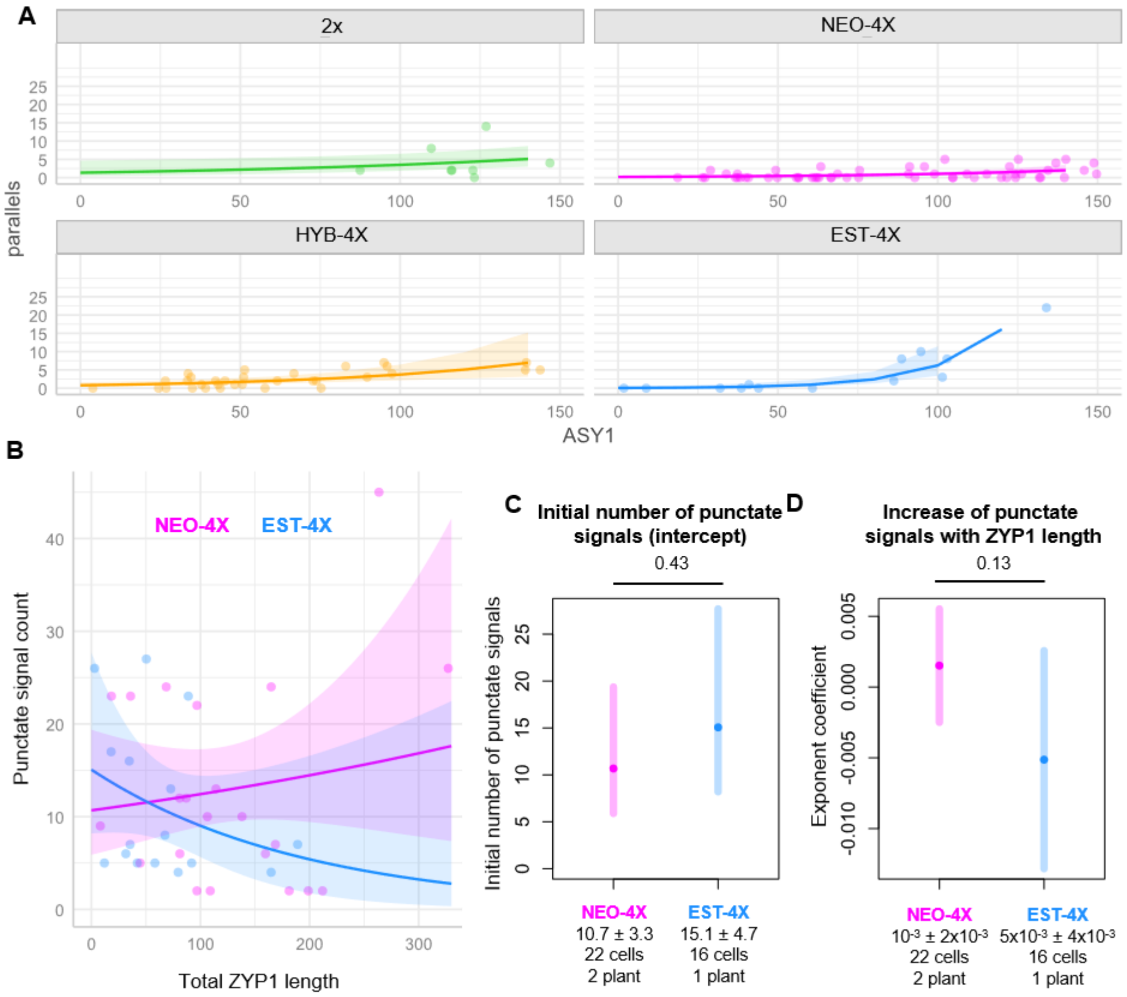


**Fig. S5.** Statistical models for synapsis defects. (A) Plots of the data and fitted trendlines predicted by a negative-binomial-GLMM explaining the change in the number of instances of parallel axes as the length of ASY1 signal increases for the four genotypes. (B) Plots of the data end the trendlines predicted by a negative binomial-GLMM for the change in the number of punctate ZYP1 signals. (C) and (D) are plots showing differences predicted by the fitted model in the intercept for both genotypes (i.e. in the initial number of punctate signals, when ZYP1 length is zero, in C) the exponent coefficient (i.e. the change of the number of punctate signals as ZYP1 length grows). P-values are indicated on top of the plots for each comparison (according to Wald's test on Poisson-GLMM coefficients). Significant p- values were highlighted in red. Mean values (Mean ± SE) and sample sizes are indicated in the lower part of the plots.

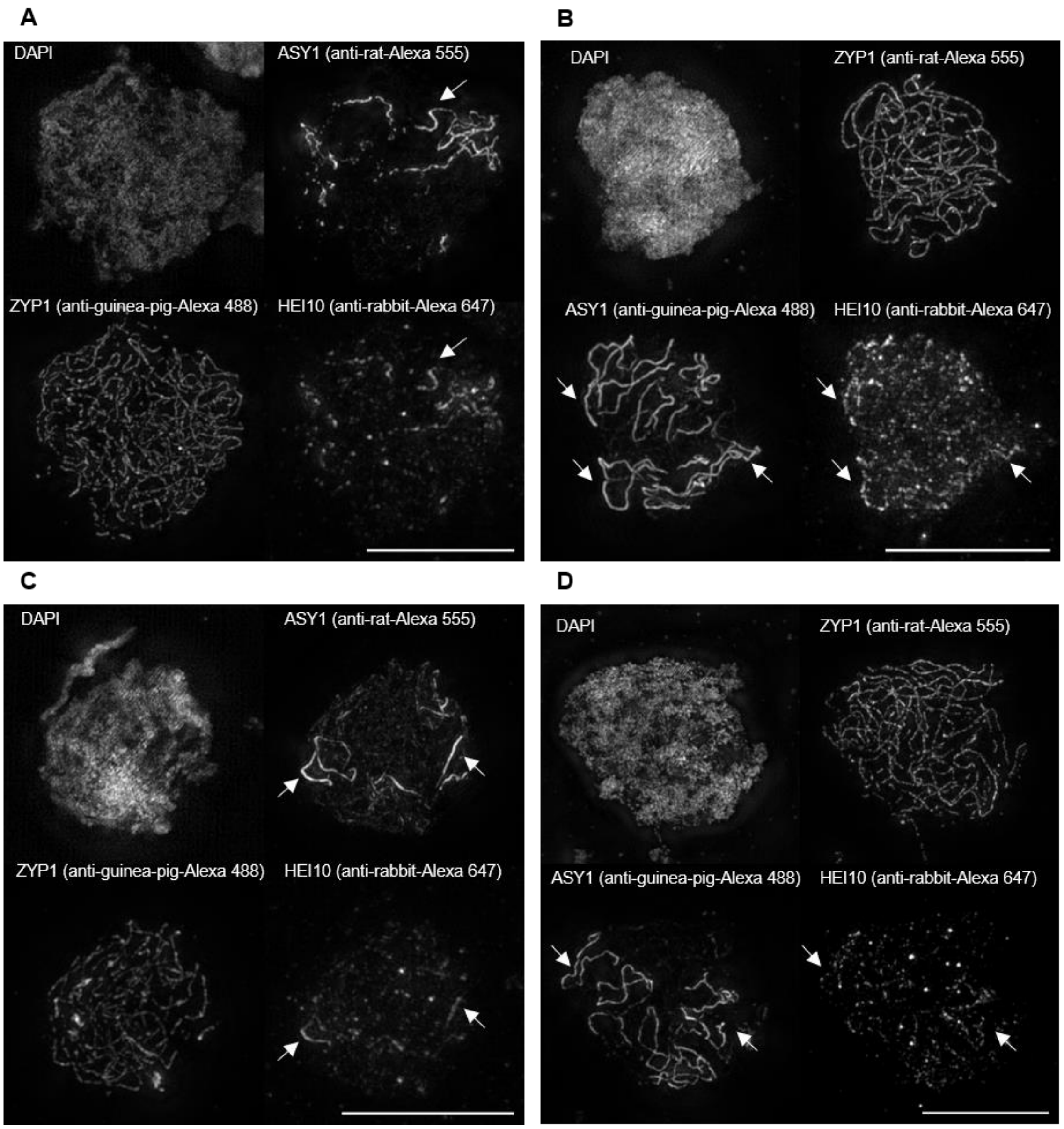


**Fig. S6.** HEI10-ASY1 overlapping revealed by different combination of antibodies. (A) and (C) show immunostaining one combination of primary and secondary antibodies whereas (B) and (D) shows a different combination.

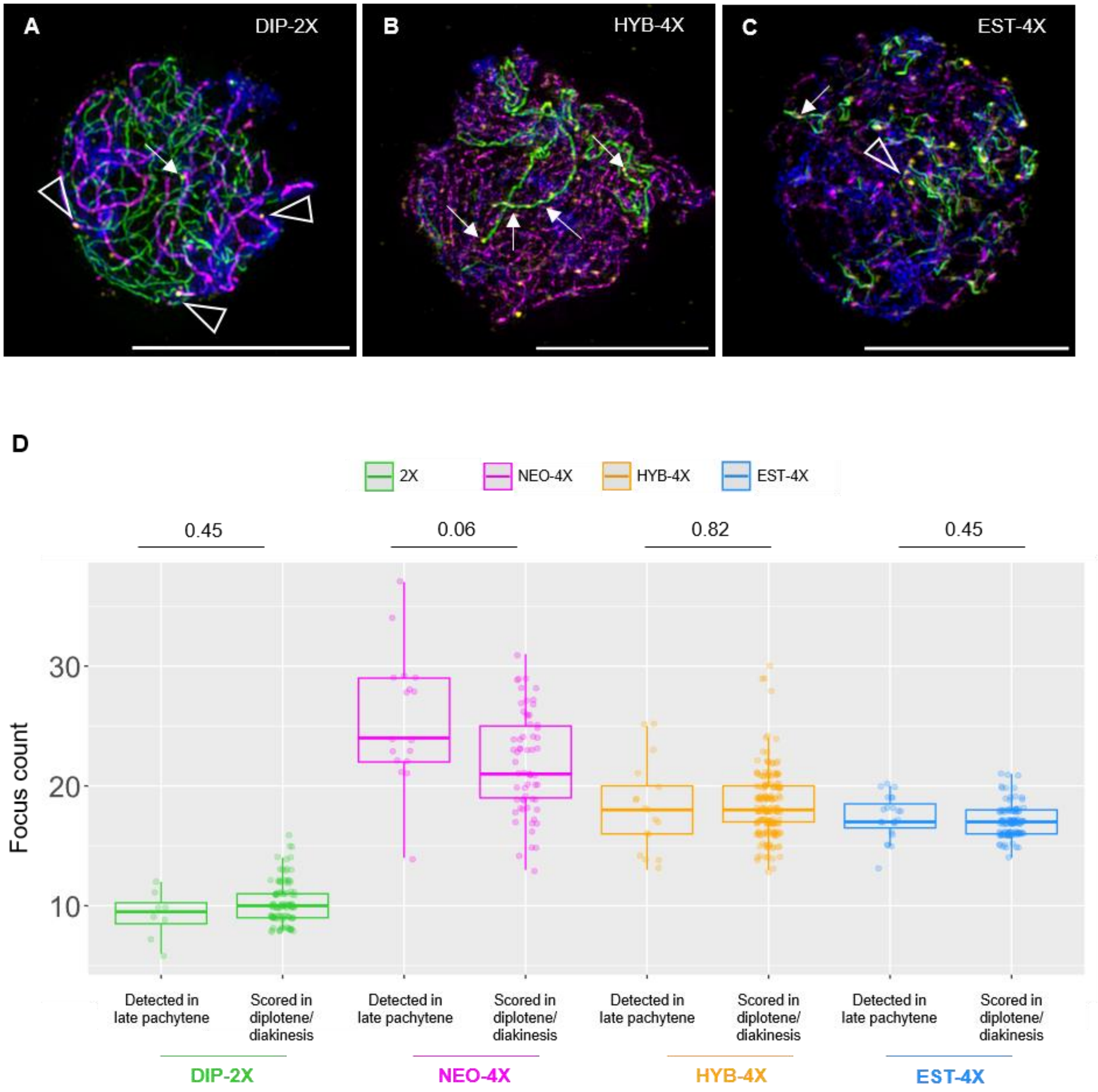


**Fig. S7.** Effect of asynapsis in HEI10 localization and crossover number. (A), (B) and (C) are examples of 2X, HYB-4X and EST-4X cells (respectively) showing limited preference of prominent HEI10 foci for synapsed regions. P-values are indicated on top of the plots for each comparison (according to T-tests, except for the EST-4X comparison which was done using Mann-Whitney U test).

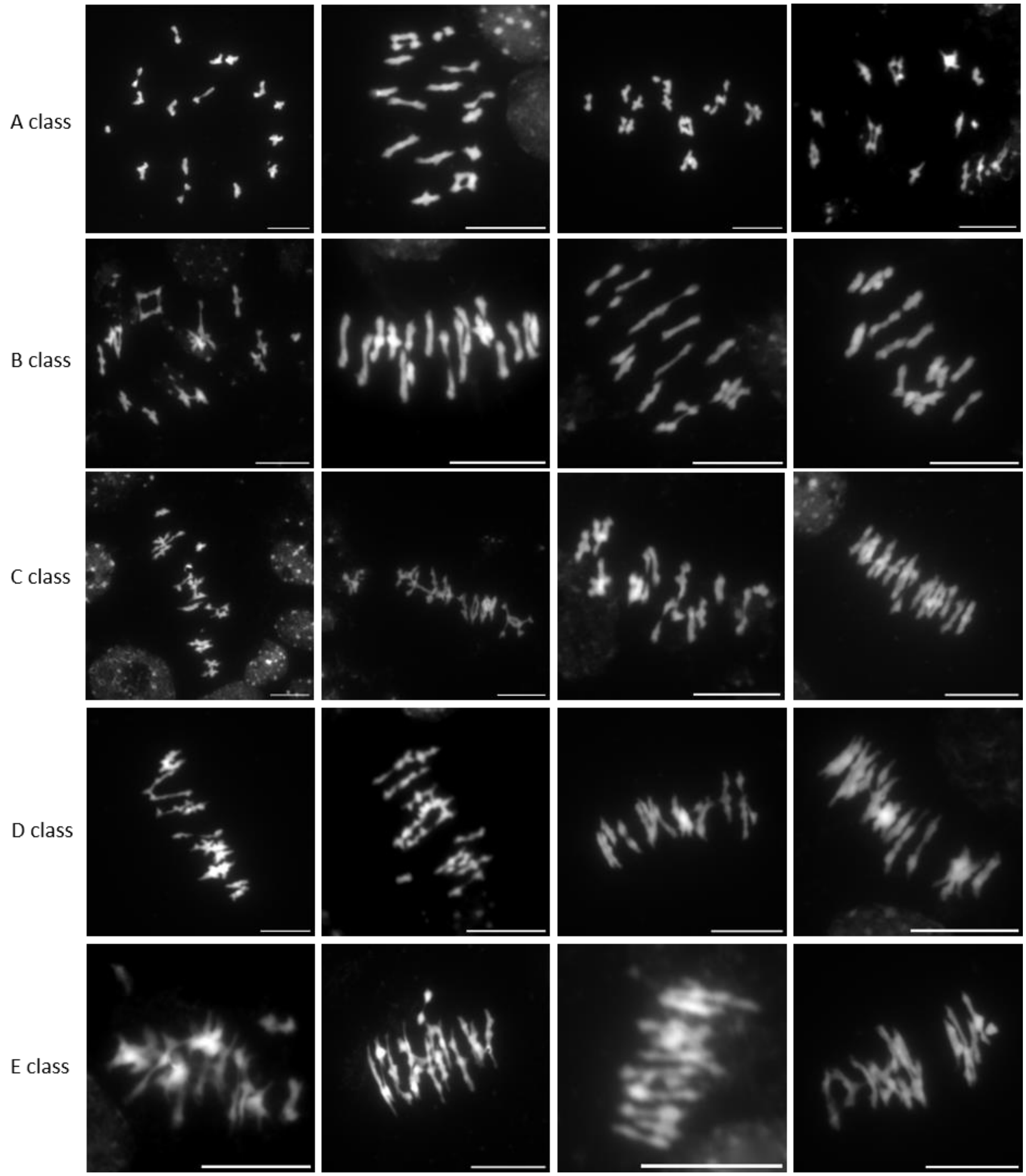


**Fig. S8.** Scorabiilty classes in metaphase I Examples of metaphase I cells assigned to different “scorability” clasess; named A, B, C, D, or E, depending on the quality of the spread and the confidence on the count, being A the gold standard and E the least reliable.

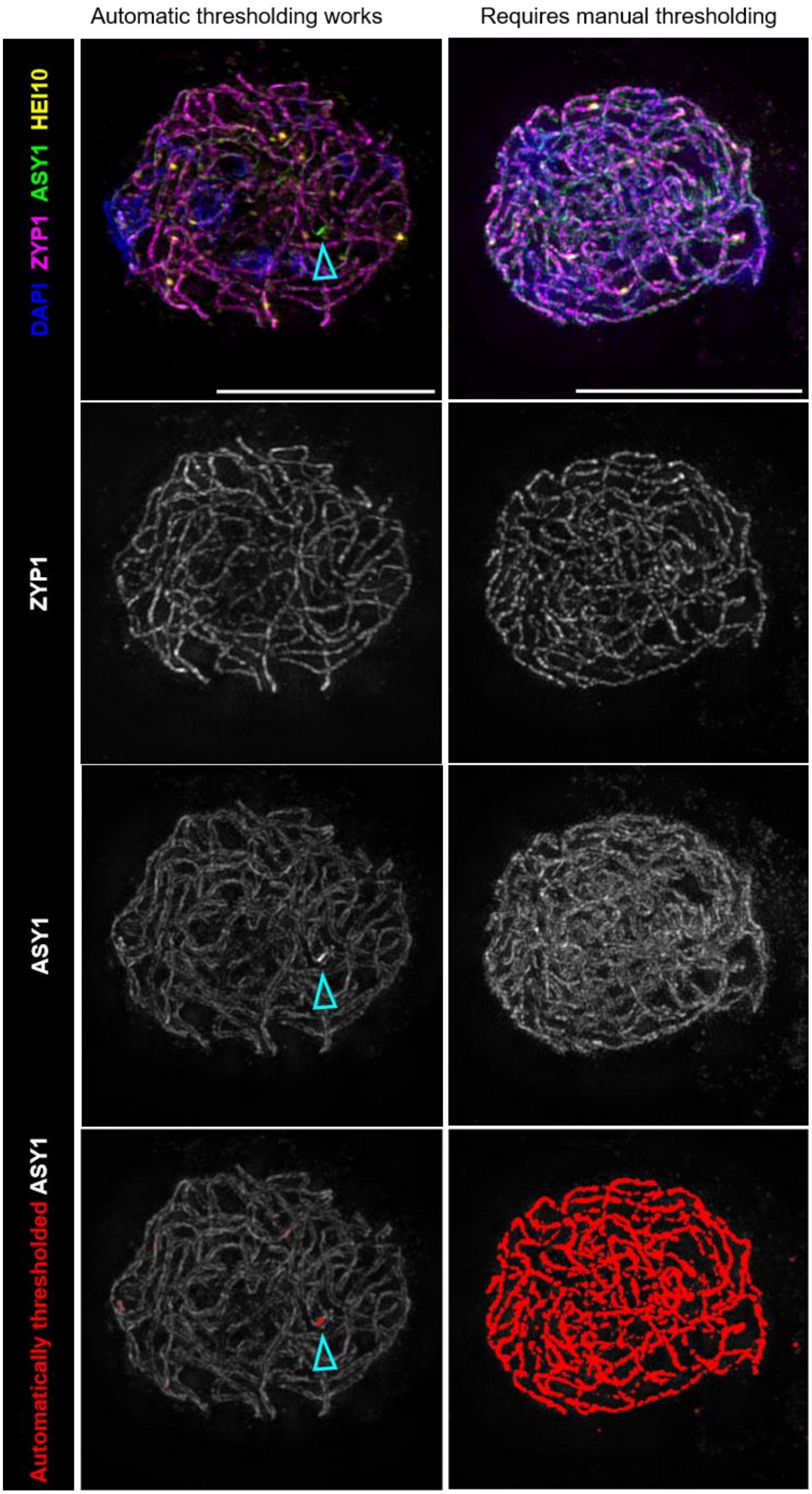

Automatic thresholding works
Requires manual thresholding
DAPI ZYP1 ASY1 HEI10
ZYP1
ASY1
Automatically thresholded ASY1

**Fig. S9**. Automatic thresholding for ASY1 signal. Examples of two whose ASY1 signals are detected differently by Yen threshold. The first cell (left) displays complete synapsis and the threshold does not detect the dim ASY1 signal characteristic of a remodeled axis after synapsis. A miniscule artifact or synaptic imperfection (cyan arrowhead), makes the threshold to detect it yielding negligible ASY1 length measurement. By contrast the other cell (right), shows also full synapsis but there is no artifact or synaptic imperfection. Therefore since the threshold operates with relative intensities (1), detects all the dim ASY1 signal. This causes a massive overestimation of the ASY1 length where massive asynapsis measurements are given for fully synapsed cell. Fortunately, these cells are rare and easily detected with Macro 3 as described in Supporting Information Text.

**Table S1.** Plant material used for this study. It is indicated whether the plant was colchicine-treated and how each plant was analyzed; either with epifluorescence microscopy (metaphase I spreads plus late prophase I HEI10 immunostaining) or SIM (for prophase I dynamics analysis).

| Plant name | Genotype | Colchicine-treated? | Use |
|---|---|---|---|
| D1-5 | 2X | No | Epifluorescence |
| D1-5 | 2X | No | SIM |
| D2-1 | 2X | No | SIM |
| SNO-1 | 2X | No | SIM |
| SNO-4 | 2X | No | SIM |
| cD4-2 | NEO-4X | Yes | SIM |
| cSN-1 | NEO-4X | Yes | SIM |
| cSN-10-1 | NEO-4X | Yes | Epifluorescence |
| cSN-2 | NEO-4X | Yes | SIM |
| N1-5 | NEO-4X | No | Epifluorescence |
| N2-1 | NEO-4X | No | Epifluorescence |
| N2-2 | NEO-4X | No | Epifluorescence |
| N2-5 | NEO-4X | No | Epifluorescence |
| N2-6 | NEO-4X | No | Epifluorescence |
| N2-9 | NEO-4X | No | Epifluorescence |
| N2-9 | NEO-4X | No | SIM |
| N3-4 | NEO-4X | No | Epifluorescence |
| N3-12 | NEO-4X | No | SIM |
| N3-13 | NEO-4X | No | Epifluorescence + SIM |
| F1 | HYB-4X | No | SIM |
| F1-2-3 | HYB-4X | No | Epifluorescence |
| F2-3-3 | HYB-4X | No | Epifluorescence |
| F3 | HYB-4X | No | SIM |
| F3-1 | HYB-4X | No | Epifluorescence |
| F3-4-1 | HYB-4X | No | Epifluorescence |
| H4-10 | HYB-4X | No | Epifluorescence |
| H4-10 | HYB-4X | No | SIM |
| H4-9 | HYB-4X | No | Epifluorescence |
| T1-2 | EST-4X | No | SIM |
| T1-20 | EST-4X | No | Epifluorescence + SIM |
| T1-23 | EST-4X | No | Epifluorescence |
| T1-5 | EST-4X | No | Epifluorescence + SIM |
| T2-1 | EST-4X | No | Epifluorescence |
| T2-10 | EST-4X | No | Epifluorescence |
| T2-3 | EST-4X | No | Epifluorescence |
| T3-1 | EST-4X | No | SIM |
